# A Highly Sensitive Fluorogenic Assay for the Detection of Nephrotoxin-Induced Oxidative Stress in Live Cells and Renal Tissue

**DOI:** 10.1101/2020.06.01.121707

**Authors:** Kamalika Mukherjee, Tak Ian Chio, Han Gu, Dan L. Sackett, Susan L. Bane, Sanja Sever

## Abstract

A common manifestation of drug toxicity is kidney injury or nephrotoxicity. Nephrotoxicity frequently leads to termination of clinical trials and drug withdrawals, which jeopardize biomedical progress. Efficient preclinical screening platforms capable of detecting mild signs of kidney damage, which may elicit considerable toxic response *in vivo,* are essential. A common manifestation of chemical toxicity is oxidative modification of cellular biomolecules. Therefore, we have developed a facile biomolecule carbonyl detection assay that well surpasses the sensitivity of the standard assays in identifying modest forms of renal injury. Using a novel fluorogenic sensor, TFCH, we have demonstrated the applicability of the assay in live kidney cells and in renal tissue. This robust assay can help inform preclinical decisions to recall unsafe drug candidates. Application of this assay in identifying and analyzing diverse pathologies is envisioned.

## 1. Introduction

Drug-induced kidney injury, or nephrotoxicity, is a critical limiting factor in the development of new therapeutics. Existing preclinical screening processes are often unable to predict nephrotoxicity in humans, which results in failed clinical trials [1,2]. Currently used phenotypic preclinical screening assays measure alterations in cell viability, morphology and mitochondrial function [2]. However, the majority of these assays detect only severe forms of injury at high doses and/or after lengthy exposure to compounds. Therefore, these assays are insensitive to modest changes that can potentially generate a much greater magnitude of toxicity *in vivo* [1]. Recently a heme oxygenase-1-based nephrotoxicity screening assay has been reported [3,4,5]. Although robust, even this cell-based assay requires high concentrations of the drugs, fixed cells and lengthy sample preparation to generate detectable signals. The need for potent screening assays for nephrotoxicity in live cells and tissue therefore remains unmet and continues to impede biomedical advancement and pose economic burden on the drug development process.

An early response to chemical toxicity is oxidative stress (OS), which is evidenced by an upsurge of oxidants, such as reactive oxygen species (ROS), and a depletion of reductants, such as reduced glutathione (GSH) [6]. While changes in the levels of ROS and GSH are transient, a cardinal irreversible consequence of OS is the carbonylation of biomolecules (Figure 1A). Since these stable modified biomolecules are formed promptly after chemical injury, they can serve as a biomarker for identifying potential cytotoxins. Carbonylation is commonly detected using alpha-effect amines as reporter molecules in biochemical assays [7,8]. The conventional assays often require lengthy tedious downstream processing and harsh chemical components that can alter subcellular structures, thereby misrepresenting spatial distribution of carbonylated biomolecules [9]. Moreover, end-point analysis of fixed cells or cell lysates was the only option until we demonstrated the first live cell compatible assay using synthetic probes, coumarin hydrazine (CH) and benzocoumarin hydrazine (BzCH) [10,11]. Our approach was also validated by Vemula *et al* using a commercially available probe, DCCH [9]. Since crucial prerequisites for identifying mild phenotypes of chemical toxicity are high sensitivity and versatility of the assay, this work is aimed at achieving these objectives. Leveraging our experience in probe development for biomolecule carbonyls, we have developed a novel fluorescent probe that is particularly suited for detecting mild signs of nephrotoxicity in live cells and living tissues.

**Fig. 1:**
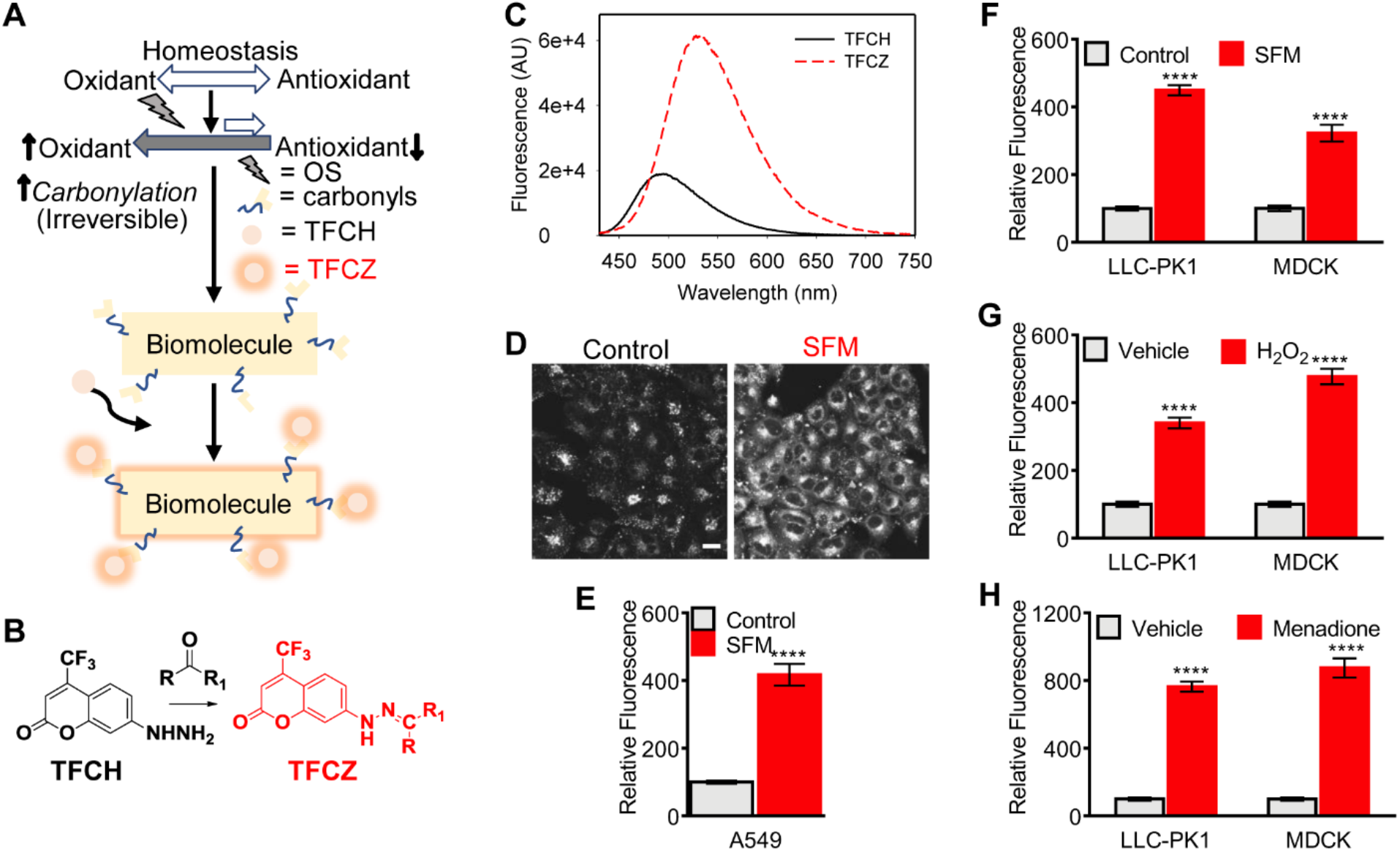
TFCH is a fluorescent probe for oxidative-stress induced carbonylation in live cells. Schematic representation of fluorogenic detection of oxidative stress-induced carbonylation (A). Chemical structure of 4-trifluoromethyl-7-hydrazinyl-2H-chromen-2-one (TFCH) and its corresponding hydrazone (TFCZ) (B). Emission spectra of 10 μM TFCH or TFCZ in phosphate buffer containing 0.5% (v/v) DMSO (C). Excitation wavelength = 405 nm. A549 cells grown in standard media (control) or serum-free media (SFM) for 24 h were allowed to react with 20 μM TFCH for 30 min, washed, and imaged live (D). Scale bar, 20 μm. Bar graphs showing quantification of cellular carbonyls detected by TFCH in live control and serum-starved live A549 cells (E). Bar graphs showing quantification of cellular carbonyls detected by TFCH in control and serum-starved LLC-PK1 or MDCK cells (F). Bar graphs showing quantification of cellular carbonyls detected by TFCH in LLC-PK1 or MDCK cells treated with 400 μM hydrogen peroxide or vehicle (water) (G). Bar graphs showing quantification of cellular carbonyls detected by TFCH in LLC-PK1 or MDCK cells treated with 100 μM menadione or vehicle (DMSO) (H). TFCH (20 μM) was added to live cells and incubated for 30-60 min, which were then rinsed, fixed and imaged (F-H). At least three independent experimental sets were performed, and fluorescence associated with > 100 cells were quantified. An unpaired t-test with Welch’s correction was performed. \*\*\*\**P* <0.0001. Error bars represent SEM.

## 2. Results and Discussion

### 2.1. TFCH is a fluorogenic sensor for oxidative-stress induced carbonylation in live cells

We have previously shown that coumarin-based fluorophores have low inherent toxicity and can be readily internalized and washed out from the cells. It was therefore desirable to retain these features in a new fluorophore. For the detection of mild chemical injury, we needed a sensor that has high sensitivity. It is known that appending a trifluoromethyl group to the coumarin scaffold strongly red shifts the absorption and emission envelopes and engenders higher photochemical stability relative to alkylated coumarins [10]. Therefore, a trifluoromethyl derivative of CH, TFCH, was designed to be a robust, live cell compatible probe to detect low levels of biomolecule carbonylation that is expected to result from mild nephrotoxicity. Using A549 lung cancer cells, we confirmed that this modification of the chromophore did not alter its live cell compatibility. TFCH was not cytotoxic and its cellular influx and efflux could be achieved seamlessly (≤ 5 min) (Figures S1F-I).

Next we prepared a hydrazone product of TFCH with an aliphatic aldehyde, TFCZ, as a model compound for carbonylation detection (Figure 1B). Hydrazone formation resulted in a bathochromic shift in the emission maximum and induced a substantial increase in the emission intensity relative to the unreacted probe (Figures 1C, S1A-E), which is a desirable photochemical property. TFCZ showed an exceptionally large Stokes shift of ~145 nm, which eliminates the chances of self-quenching and is a generally useful feature for analytical assays (Figures S1). Since high sensitivity of the assay is an essential prerequisite, TFCZ was used to determine the assay sensitivity in a biological milieu. The limit of detection (LOD) of TFCZ was ~89 nM in cell lysate (of A549 lung cancer cells), which is an order of magnitude smaller than that of our previously reported fluorophore [11]. These data strongly support the idea that TFCH can serve as a highly sensitive and photochemically desirable reporter molecule for mild signs of chemical toxicity.

Different cell-based assay formats with TFCH were then explored. Using serum-free media (SFM) as an OS-induction model, we showed that TFCH can be used in a simple platereader-based assay suitable for high throughput screening (HTS) (Figure S2A). Importantly, it was possible to identify carbonyls in live cells in the presence of unreacted fluorophore by a one-step high content screening (HCS)-compatible assay (Figure S2B). This single-step (no wash) assay format may be particularly suitable for screening OS-inducing molecules that have a propensity to induce cell detachment. The high sensitivity of the TFCH assay was further validated by the fact that only 2 μM of TFCH was necessary to detect serum starvation-induced carbonylation in live cells. This is 1/10^th^ of the fluorophore concentration needed for CH and BzCH-based assays and 1/400^th^ of the fluorophore concentration reported for the DCCH assay [9,10,11].

Other formats of the assay can also be used. For example, excess fluorophore can be removed and the cells can be either imaged live (Figures 1D,1E) or can be fixed and preserved for analysis at a later time (Figure S2C). Indeed, a similar spatial distribution and increase in carbonylation due to serum starvation was detected in both live (Figures 1D, S2B) and fixed cells (fixed after fluorescent labeling in live cells) (Figure S2C). Together, these data establish TFCH as a highly sensitive tool for visualizing and quantifying biomolecule carbonyls in live cells using multiple assay formats.

### 2.2. A sensitive fluorescent assay for screening chemical toxin-induced OS in live renal cells

To assess the potential of using TFCH for detecting nephrotoxins, we chose to focus on two standard cellular models for assessing small molecule toxicity: porcine kidney proximal tubule (LLC-PK1) cells and distal tubule-derived Madin-Darby Canine Kidney (MDCK) cells [12,13]. We established our assay in these cell lines using SFM, menadione or hydrogen peroxide (H_2_O_2_) as stress induction models. TFCH detected significant increase (~220 to ~770%) in carbonylation in both cells lines subjected to the aforementioned OS-inducing agents. (Figures 1F-H and S2D-F).

We used two drugs with known nephrotoxicity in humans, cisplatin (anticancer) and gentamicin (antibiotic), to demonstrate and validate our tool and assay [14]. The high expression level of copper transporters responsible for cisplatin endocytosis in proximal tubule cells makes these cells more vulnerable to the cisplatin injury [15,16]. The same cells are also the primary site of injury for gentamicin, which is endocytosed by megalin and cubilin complex [17]. Loss of cell polarity of these renal epithelial cells and alteration in the actin cytoskeleton are prominent manifestations of nephrotoxicity [18,19,20]. In addition, both drugs increase OS and initiate cell signaling pathways that ultimately lead to cell death and/or detachment [21,22]. The aforementioned phenotypes, including change in cell morphology, status of the OS and cell viability are the basis of conventional nephrotoxicity screenings. Therefore, in validating our assay for nephrotoxicity, we examined how it compares with other available assays.

Since our goal was to develop an assay that can detect early and modest injury, we first focused on determining experimental parameters that produce only sub-cytotoxic effects after drug treatment. To facilitate the detection of early signs of drug toxicity, we analyzed the cells after 24 h drug exposure, instead of the usual 72 h [2]. To achieve our second objective of detecting mild injury, we chose drug concentrations that showed minimal toxicity (cell viability ≥ 65%) based on a resazurin assay (monitors cell metabolism) and a sulforhodamine B (SRB) assay (monitors cell number) (Figure 2).

**Figure. 2:**
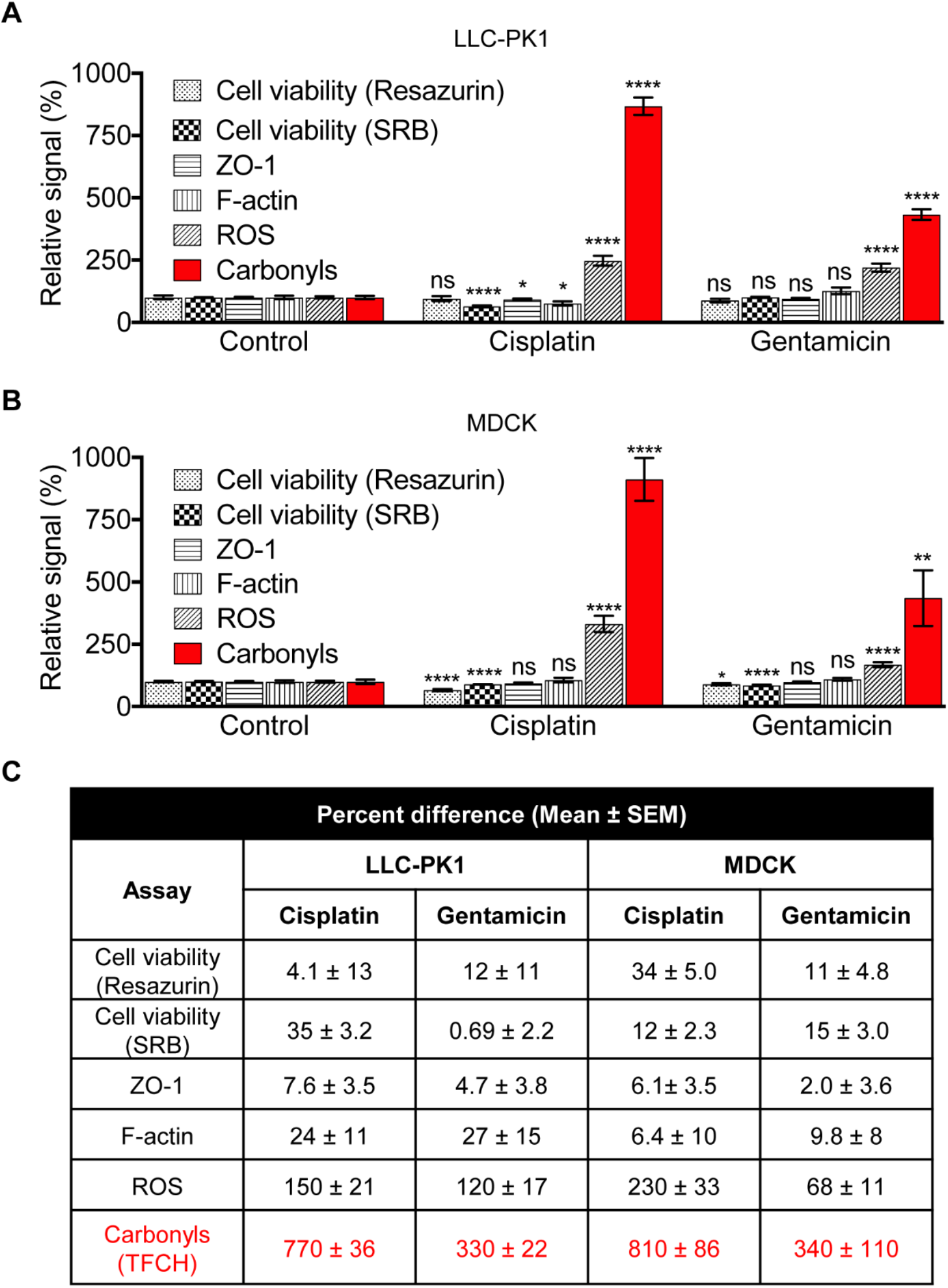
TFCH-mediated detection of carbonylation outperforms classical nephrotoxicity assays in renal epithelial cells. Bar graphs showing the effects of cisplatin (1.5 μg/mL) or gentamicin (0.58 mg/mL) on LLC-PK1 (A) or MDCK (B) cells after 24 h. Cell viability was assessed by a resazurin or SRB assay (independent experimental sets, n ≥ 2). Level of ZO-1 or actin stress fibers (F-actin) (n ≥ 2, number of cells quantified per condition, cell no. ≥ 60) was evaluated by immunocytochemistry; reactive oxygen species (ROS) or carbonylation (n ≥ 4, cell no. ≥ 200) level was assessed by CellROX Green or TFCH respectively; as described in *Methods*. Error bars represent SEM. Percent difference of each treatment from the control (no drug treatment) recorded by each assay (C). An unpaired t-test with Welch’s correction was performed. \*\*\*\**P* <0.0001, \*\*\**P* <0.001, \*\**P* <0.01, \**P* <0.05, *P*>0.05 was considered not significant (ns).

We next assessed structural damage in cells under identical conditions by examining the extent of loss of cell polarity and alterations in the actin cytoskeleton, two common phenotypes of nephrotoxicity. We quantified the intensity of zonula occludens-1 (ZO1), a tight junction protein that defines cell polarity [23,24], and the level of intracellular actin stress fibers (represented by filamentous actin, F-actin). No change in cell morphology was observed, except for a minor difference in LLC-PK1 cells with cisplatin (Figures 2A,B). Overall, these assays were deemed not sufficiently sensitive to detect cellular damage under these conditions.

We next examined the effects of the drugs on the status of cellular OS by determining the level of ROS, a dominant precursor of carbonylated biomolecules. Although we established the assay conditions to generate mild cellular phenotypes, a significant increase (~68% – 230%) in the level of ROS was observed in both cell lines treated with gentamicin and cisplatin. (Figure 2, S3). These data show that the ROS levels exhibit the greatest magnitude of difference between injured and uninjured cells, suggesting that OS is the strongest and earliest phenotype tested thus far.

Finally, we tested TFCH’s ability to detect drug-induced cellular carbonyls, a stable downstream effector of ROS, under the same conditions. TFCH showed a strong response with both drugs in both cell lines (Figure 2, S3). The injured cells were ~300 – 800% more fluorescent than their uninjured counterpart when assayed with TFCH, whereas the best signal enhancement from the ROS assay was < 250 %. A side-by-side comparison of the changes in cell morphology or OS level generated by subtoxic concentrations of cisplatin and gentamicin clearly illustrates the superiority of TFCH assay (Figure 2C).

Owing to the early induction of the biomarker and the desirable sensitivity of the fluorophore, we speculated that our assay may be able to detect injury after a brief exposure to the drugs, instead of the usual 24 h. We treated the cells with the drugs for 3 h prior to adding TFCH. We observed a significant increase in fluorescence, up to ~190%, in cisplatin-or gentamicin-injured cells (Figure S2G-I). The temporal sensitivity, along with the ease of performing the assay, enable data generation within hours instead of days. Together, these data show that the TFCH assay is demonstrably the most sensitive assay tested herein and is capable of measuring early signs of drug insult.

### 2.3. TFCH is a probe for detecting carbonylation in live renal tissue

While monolayers of renal cells in culture serve as the current gold standard for screening nephrotoxins, the complexity of the renal system as a whole is not comprehensively represented by any single cell type in culture [1,25]. More physiologically relevant screening platforms are critical for improving safety profile predictability. We thus tested the applicability of TFCH in detecting kidney tissue injury in live tissue slices. The kidney slices were maintained live during the experimental procedure using a pre-established protocol to support normal physiology of the tissue [26]. Rat kidney slices subjected to cisplatin or gentamicin treatment followed by TFCH exhibited substantially higher levels of carbonyls compared to the uninjured control (Figures 3, S4). The fluorescent labeling was mainly associated with the tubules and not the glomerulus (Figure S4B), which is in agreement with the existing paradigm that renal proximal tubules are the primary sites of drug-induced damage [16,17,27]. Our data thus attest to the utility of this assay and fluorophore in a complex tissue system.

**Fig. 3:**
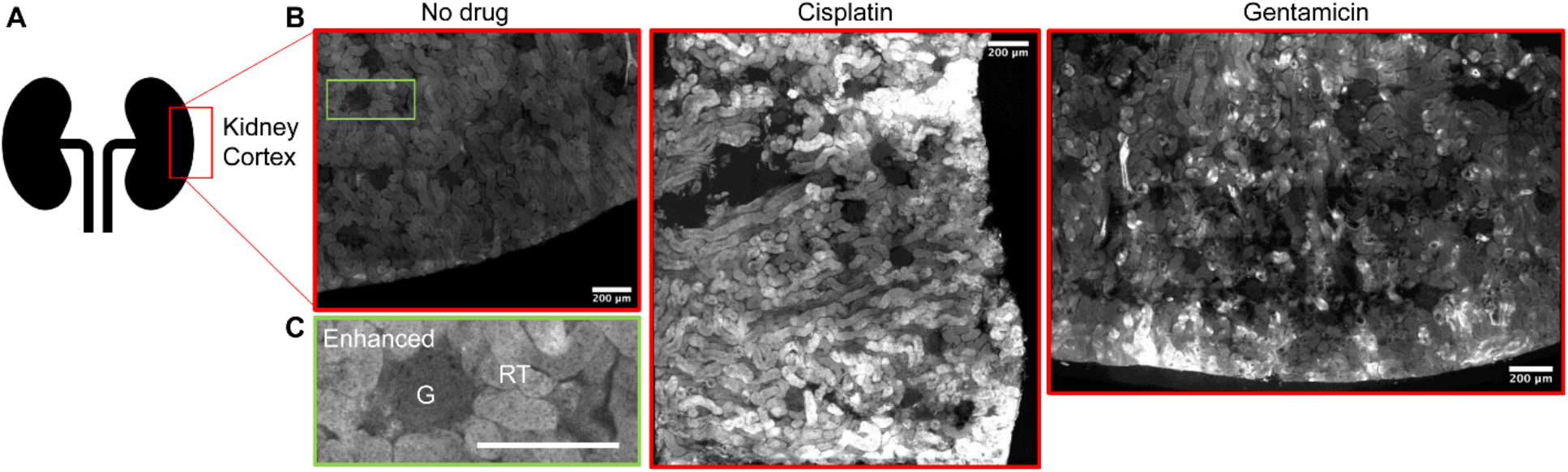
TFCH detects drug-induced carbonylation in live rat kidney slices. Schematic representation showing the region of kidney used for imaging (A). Representative photomicrographs assembled (stitched) from multiple sections of the renal cortex (B). As indicated, rat kidney slices were exposed to cisplatin (150 μg/mL), gentamicin (4.6 mg/mL), or vehicle (buffer; no drug) for 1 h, followed by addition of TFCH (2 *μ*M) for 30 min. An inset (enhanced) (C) of the control (no drug) slice is showing the location of renal tubule (RT) and glomerulus (G). Scale bar, 200 μm

In summary, we have developed a rapid and sensitive assay that probes for an early biomarker (OS-induced biomolecule-carbonyls) that is common to drug-induced nephrotoxicity. Photochemical attributes of our fluorophore and the appropriateness of the biomarker allow for significantly improved sensitivity when compared to currently used assays. The ability to detect early signs of nephrotoxicity in kidney epithelial cells by using HCS/HTS platforms is expected to facilitate facile preclinical drug safety screening [28].

Given the diversity of kidney injury, but the commonality in cellular response, fluorescent detection of carbonyls by TFCH can potentially be used for a multitude of renal injury models. In conjunction with classical histology, this assay may serve as a reliable component of a composite scoring system accounting for both structural changes (classical histology) and chemical changes (carbonylation) associated with various tissue injury models. Additionally, since the appearance of biomolecule-associated carbonyls precedes structural changes identified by classical histology, the assay presented here is expected to read early signs of modest tissue injury. The prevalence of the biomarker and the adaptability of the assay to both cell monolayer and a tissue system support the notion that insult to other organs, not confined to the kidney, can be probed by TFCH. The applicability of the assay can presumably be extended to disease models such as cancer and neurodegenerative disorders. Furthermore, the versatility of the assay should enable its application in evaluating a wide variety of pathologies and drug-induced tissue damage.

## Acknowledgement

This work was supported in part by the NIH (Grant R15 GM102867 and R15 CA227747 to SLB; R01 DK093773 and DK087985 to SS), and in part by the Intramural Research Program of the Eunice Kennedy Shriver National Institute of Child Health and Human Development (DLS). The Regional NMR Facility at Binghamton University is supported by the NSF (CHE-0922815). The authors thank Dr. Anthony Sorrentino for his initial efforts in the synthesis of TFCH and its derivative, David Tuttle for his expert assistance in image processing, Prof. Ming An for the generous gift of A549 cells, Bradley Pedro for technical assistance with cell culture, immunocytochemistry and image quantification, Prof. Richard Bouley for kindly providing the rat kidney slices and the essential facility to perform tissue-based experiments and Dr. Anilkumar Nair for his expert assistance and training in tissue imaging. Some imaging was performed in the Microscopy Core of the Program in Membrane Biology, which is partially supported by the Centre for the Study of Inflammatory Bowel Disease Grant DK043351 and the Boston Area Diabetes and Endocrinology Research Center (BADERC) Award DK057521. One of the Zeiss confocal systems was purchased with an NIH shared instrumentation grant 1S10OD021577-01.

## Author contributions

SLB and SS were responsible for acquisition of the financial support. KM and SB conceptualized the project. KM, SLB and SS designed the research in consultation with DLS. KM wrote the paper in consultation with TIC. TIC, DLS, SLB and SS edited the paper. KM, TIC and HG designed and performed the experiments, and analyzed the data.

