## Supplementary Material for "A Highly Sensitive Fluorogenic Assay for the Detection of Nephrotoxin-Induced Oxidative Stress in Live Cells and Renal Tissue"

Supplementary Figures
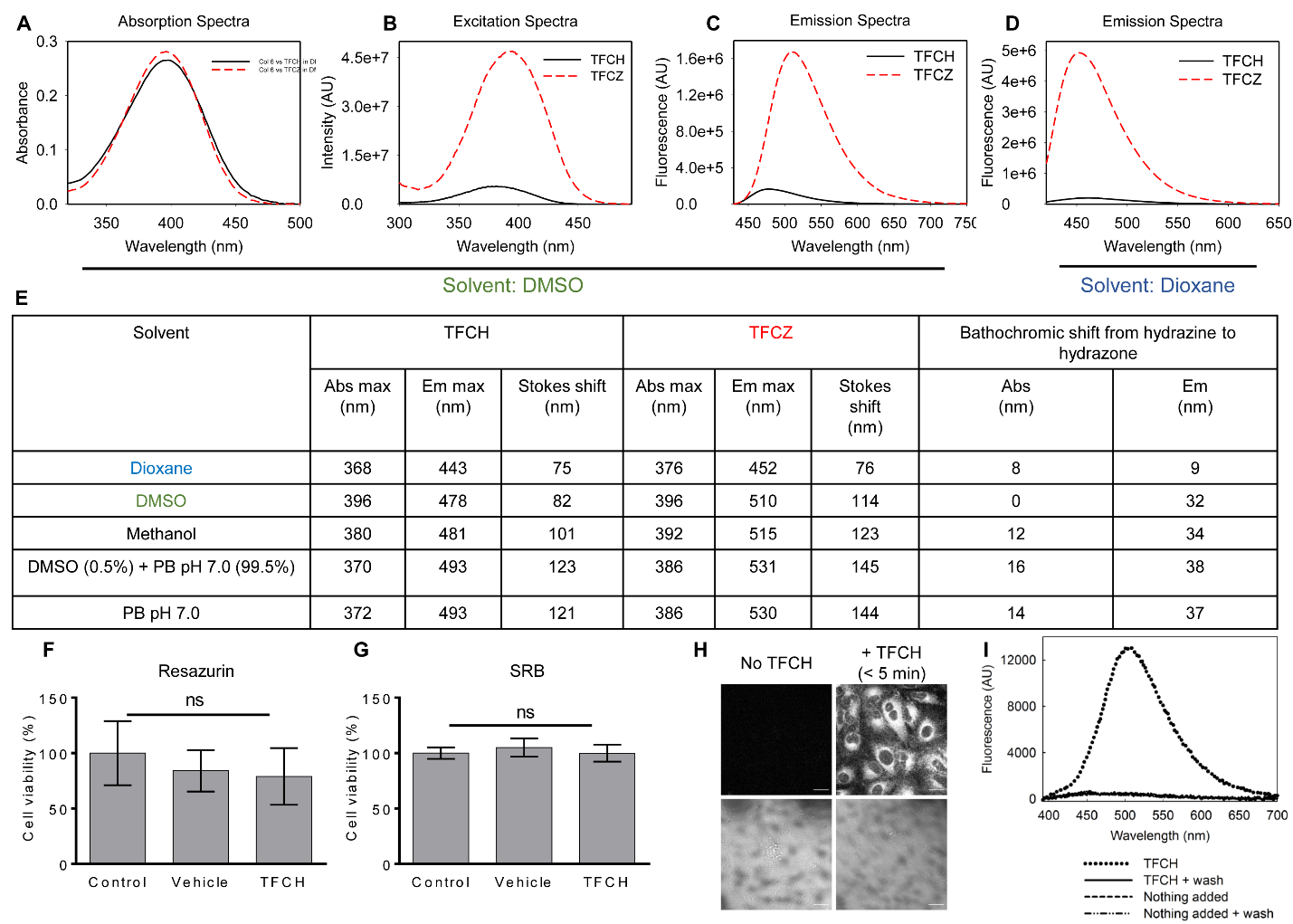

**Figure S1. TFCH is a live cell compatible probe with desirable photochemical properties.**

*Photochemical properties of TFCH and TFCZ (A-E)*. Absorption (A)**,** excitation (B), emission (C) spectra of 10$\mu$M TFCH and TFCZ in DMSO. Emission spectra of TFCH and TFCZ in dioxane (D). Spectroscopic (absorption and emission) characterization of TFCH and TFCZ in different solvents (E).

*TFCH is a live-cell compatible tool for detecting oxidative stress-induced carbonylation (F-I).* Resazurin (F) or SRB (G) assay was performed after MDCK cells were exposed to media without added treatment (control), with vehicle (0.5%, v/v, DMSO) or with TFCH (20 $\mu$M, the highest concentration used) for 24 h. Error bars represent SD. One-way ANOVA with Dunnett’s multiple comparisons test was performed to compare each treatment with the control; *P*>0.05 was considered not significant (ns). Cellular influx of TFCH occurs seamlessly (H). Live A549 cells were imaged before (no TFCH) and < 5 min after adding TFCH (+TFCH) to the cells. Scale bar, 20 $\mu$m. TFCH can be effectively washed out of live A549 cells (I). The cells were incubated with 20 $\mu M$ TFCH for 5 min. They were then either lysed immediately, or washed and lysed. Emission spectra of the lysates were recorded by exciting the samples at 372 nm.

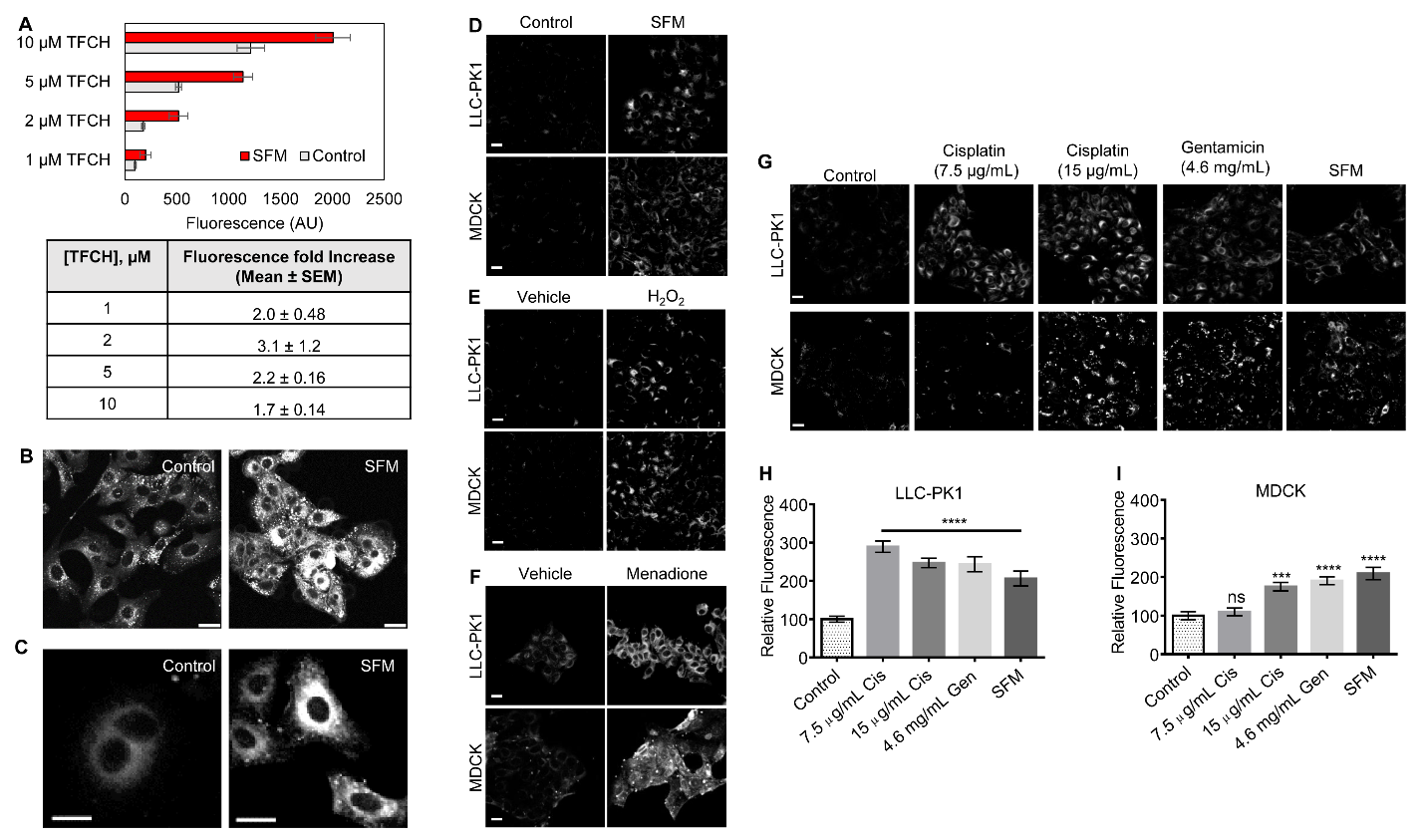

**Figure S2. TFCH can detect carbonylation induced by different models of OS in multiple cell types.**

*TFCH serves as a tool for detecting oxidative stress-induced carbonylation in live cells by HCS- and HTS-compatible assays (A-C)*. A plate-reader based assay for detecting SFM-induced carbonylation (A). A549 cells were grown in standard media (control) or SFM for 24 h before adding the stated concentration of TFCH for 90 min. The cells were rinsed with PBS and the emission was recorded at 525 nm (excitation: 405 nm). Graph showing relative fluorescence. The fold-increase in fluorescence due to serum starvation. Error bars represent SEM. A one-step assay for visualizing biomolecule-carbonyls in live cells. Scale bar, 20 μm (B). Serum-starved or control cells were allowed to react with 2 $\mu$M TFCH for 30 min before imaging the live cells without rinsing excess fluorophore. A549 cells grown in standard media (control) or SFM for 24 h were allowed to react with 20 µM TFCH for 60 min, washed, fixed, and imaged (C). Scale bar, 20 $\mu$m.

*TFCH detects oxidative stress-induced carbonylation in kidney-specific cells (D-F)*. SFM (D)-, H_2_O_2_ (E)-or menadione (F)-induced carbonylation was detected by incubating live LLC-PK1 or MDCK cells with 20 µM TFCH for 30-60 min. Cells were rinsed with PBS and fixed before imaging. Scale bar, 20 µm.

*Drug-induced carbonylation can be detected within hours by TFCH (G-I)*. LLC-PK1 (G,H) and MDCK (G,I) cells were allowed to grow for 3 h in the presence of cisplatin (Cis) or gentamicin (Gen). TFCH (20 µM) was added to the medium for an additional 45-60 min. The cells were then rinsed with PBS, fixed and imaged. One-way ANOVA with Dunnett’s multiple comparisons test was performed to compare each treatment with the control. *****P* <*0.0001*, ****P<0.001*. Error bars represent SEM. Scale bar, 20 µm.

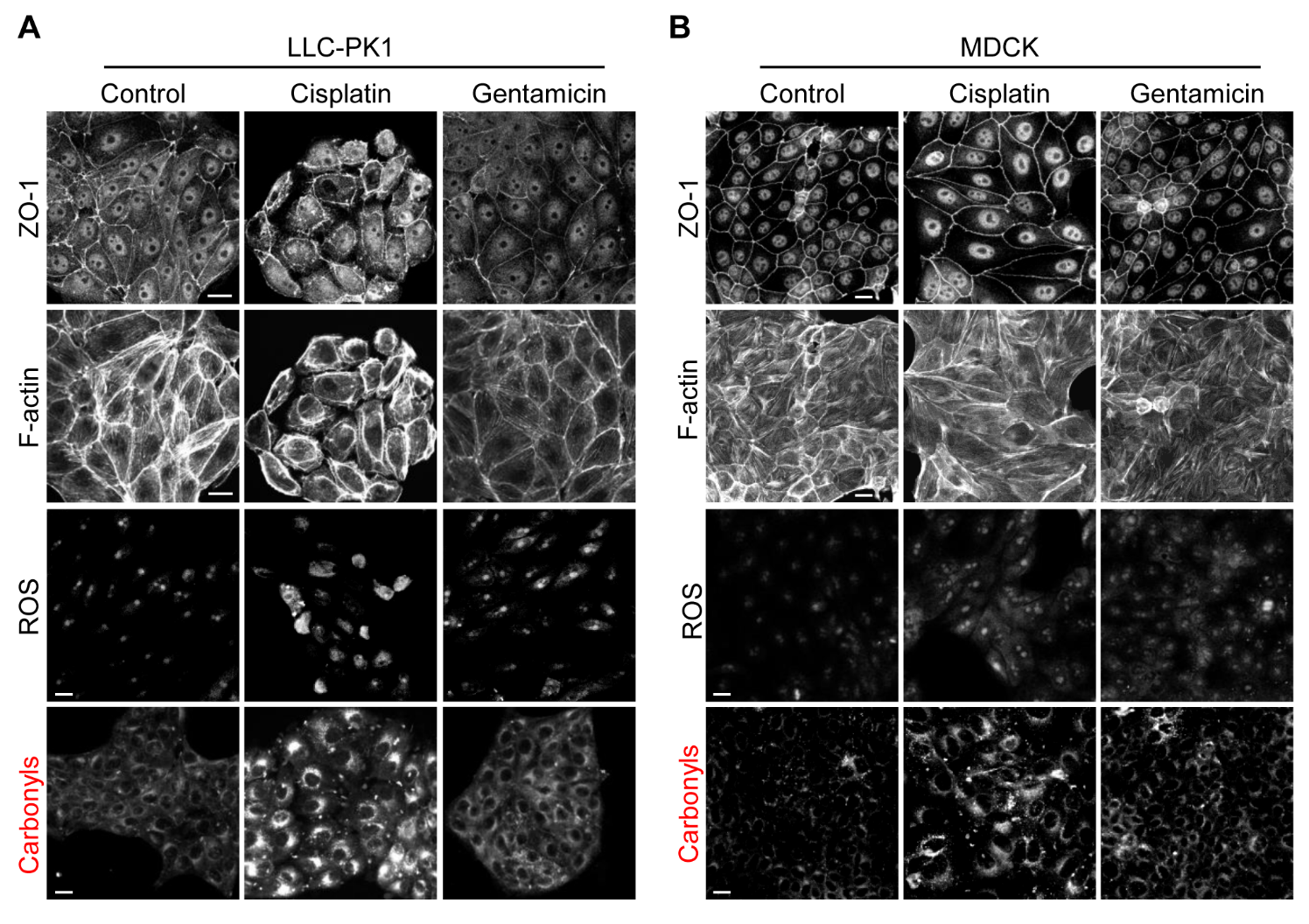

**Figure S3: Photomicrographs showing the effect of cisplatin and gentamicin in renal epithelial cells.** Status of ZO-1, F-actin, ROS, and biomolecule-carbonyls (detected by TFCH) in LLC-PK1 (A) and MDCK (B) cells. Renal cells were treated with vehicle (cell culture media; no drug; control) cisplatin (1.5 µg/mL) or gentamicin (0.58 mg/mL) for 24 h before performing the assays as described in *Methods*. Scale bar, 20 µm.

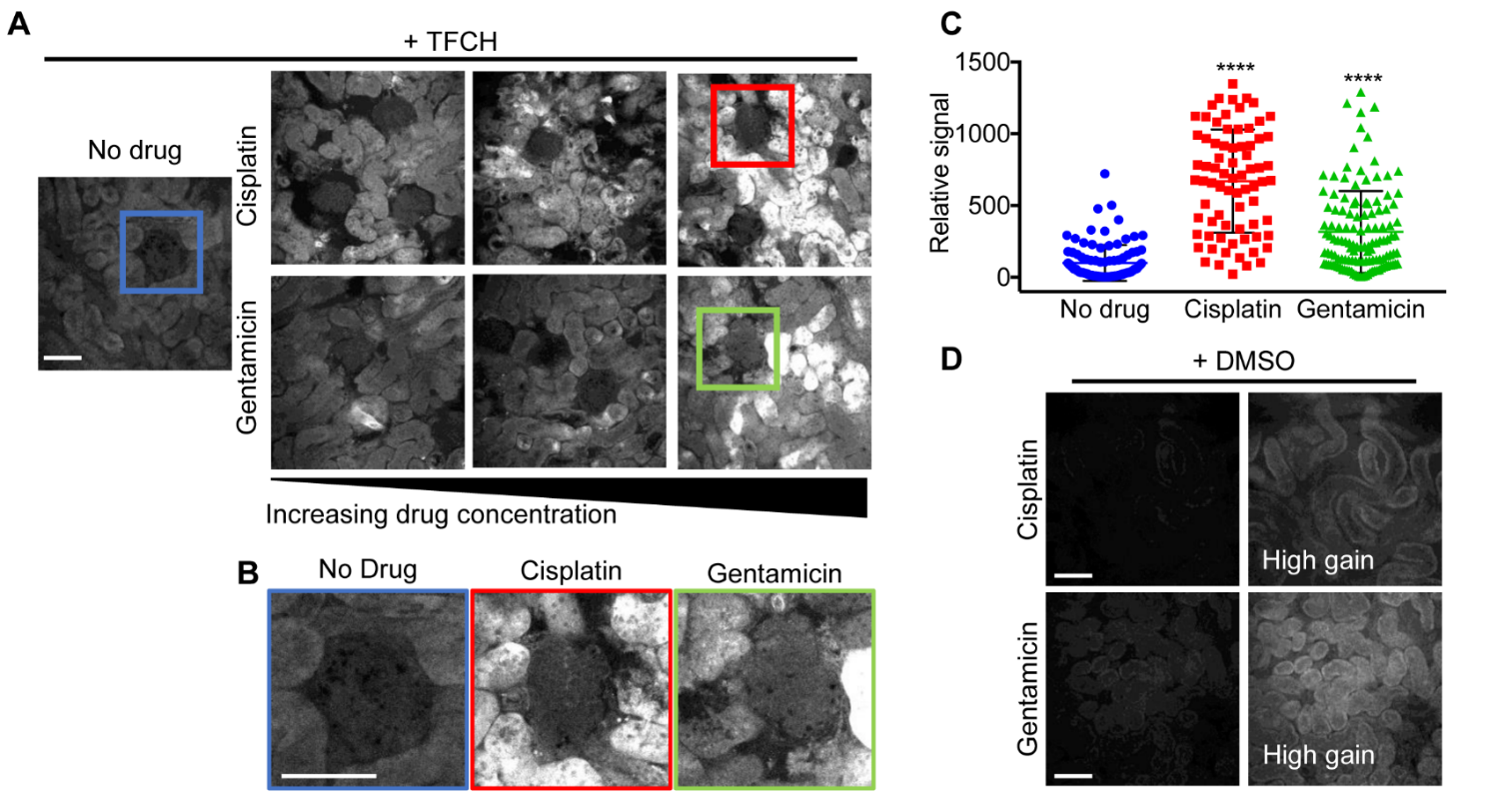

**Figure S4: Drug-induced oxidative injury detected in kidney tissue slices by TFCH.** Dose-dependent increase in carbonylation detected by TFCH (2 $\mu$M) (A). Cisplatin concentrations used: 37.5, 75, 150 µg/mL; gentamicin concentrations used: 1.2, 2.3, 6.4 mg/mL. Insets of (A) showing distribution (glomerular versus renal tubule) of biomolecule-carbonyls (fluorescence) in the tissue section (B). Quantification of renal tubule associated fluorescence in control (no drug), cisplatin (150 µg/mL), or gentamicin (4.6 mg/mL) treated tissue slices (C). Error bars represent SD. Control samples (D): tissue sections treated with cisplatin (150 µg/mL) or gentamicin (4.6 mg/mL) were exposed to vehicle (0.5% DMSO, v/v), instead of TFCH. An unpaired t-test with Welch’s correction was performed to compare the control with each treatment. *****P* <0.0001. Error bars represent SD. Scale bar, 100 µm.

**Methods**

**Synthesis and characterization of TFCH and TFCZ**

4-Trifluoromethyl-7-hydrazinyl-2H-chromen-2-one (TFCH)

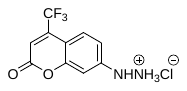

7-Amino-4-(trifluoromethyl) coumarin (500 mg, 2.18 mmol) was dissolved in 1.50 mL concentrated HCl and stirred for 10 min at -10 ºC. A chilled solution of sodium nitrite (181 mg, 1.2 eq) in 600 μL water was added dropwise to keep the temperature of reaction below 0 ºC. The solution was stirred for 1 h at -10 ºC. Stannous chloride dihydrate (1.57 mg, 3.8 eq) was dissolved in 1.50 mL concentrated HCl and chilled on ice. This cold stannous chloride HCl solution was then slowly added to the diazonium solution and the temperature of reaction was maintained below 0 ºC. The reaction was stirred for 1.5 h at -10 ºC. The yellow slurry was then filtered and washed with cold water and cold ethanol. TFCH HCl salt was collected as a light yellow solid (350 mg, 57% yield).

**^1^H NMR** (400 MHz, DMSO-d6) δ: 10.49 (br, **NH**, 3H), 9.32 (s, **NH**, 1H), 7.59 (d, J = 8.0 Hz, 1H), 7.00 (m, 2H), 6.77 (s, 1H).

**^13^C NMR** (400 MHz, DMSO-d6) δ: 159.25, 155.98, 151.00, 126.06, 123.59, 120.86, 112.53 (q, CF_3_), 111.87, 105.97, 100.31.

**MS-ESI^+^:** C_10_H_7_F_3_N_2_O_2_ [M + H]^+^ calcd.: 245.17, found: 245.12.

7-(2-Propylidenehydrazinyl)-4-(trifluoromethyl)-2H-chromen-2-one (TFCZ)

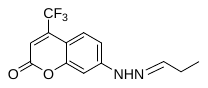

The TFCH HCl salt (20 mg, 0.082 mmol) was dissolved in 600 μL methanol and 20 μL of trifluoroacetic acid. Propionaldehyde (58.7 μL, 10 eq) was added to the solution which was stirred further for 30 min. The yellow precipitate was filtered and washed with cold methanol. The TFCZ was dried by air and collected as yellow solid (15 mg 64% yield).

**^1^H NMR** (400 MHz, DMSO-d6) δ: 10.66 (s, **NH**, 1H), 7.50 (dd, J = 9.0, 2.1 Hz, 1H), 7.37 (t, J = 4.9 Hz, 1H), 6.94 (dd, J = 9.0, 1.7 Hz, 1H), 6.86 (d, J = 2.0 Hz, 1H), 6.57 (s, 1H), 2.30 (dq, J = 7.5, 5.0 Hz, 2H), 1.08 (t, J = 7.4 Hz, 3H).

**^13^C NMR** (400 MHz, DMSO-d6) δ: 159.65, 156.80, 150.37, 147.22, 126.43, 123.75, 121.01, 110.27, 109.71 (q, CF3), 104.11, 97.76, 25.76, 11.21.

**MS-ESI^+^:** C_13_H_11_F_3_N_2_O_2_ [M + H]^+^ calcd.: 285.08, found: 285.11.

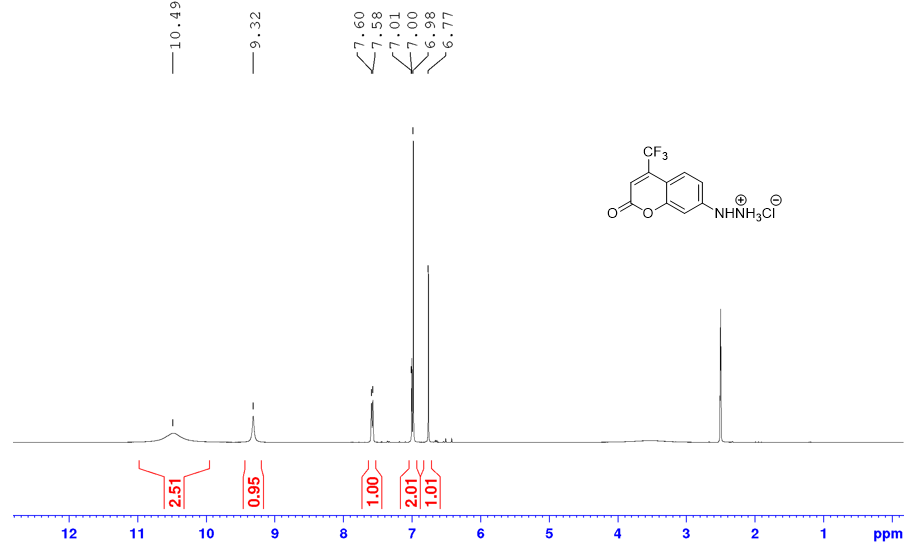

1H NMR spectrum of TFCH.

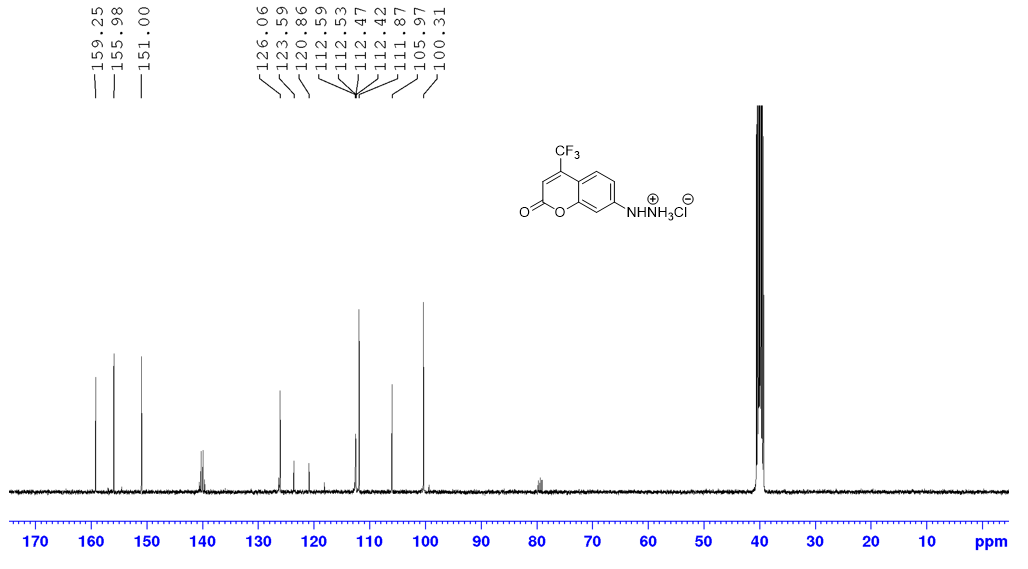

13C NMR spectrum of TFCH.

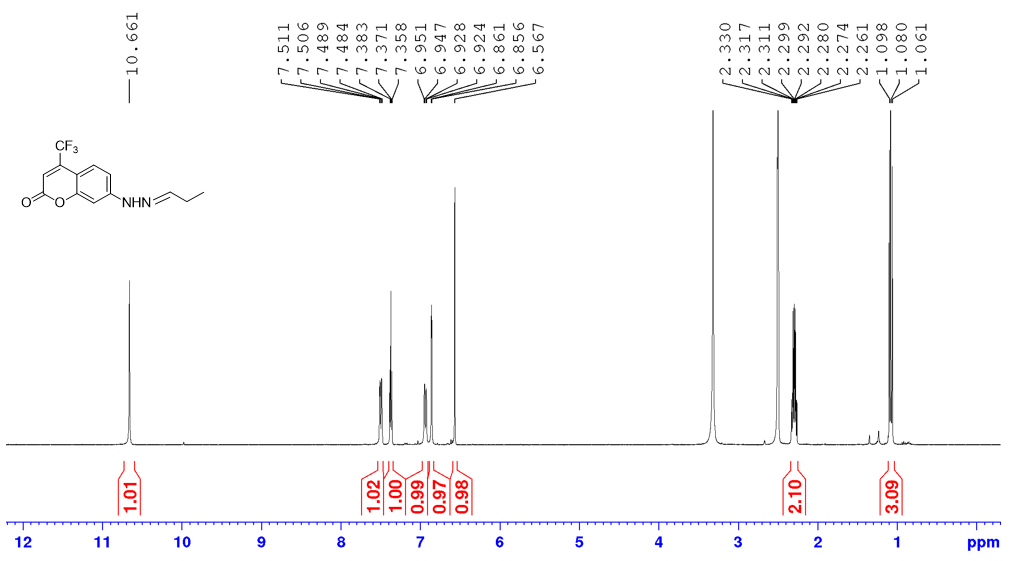

1H NMR spectrum of TFCZ (hydrazone product of TFCH and propionaldehyde).

**Spectroscopic characterization of TFCH or TFCZ**

~10 μM TFCH or TFCZ (hydrazone of TFCH and propionaldehyde) was prepared in DMSO or other solvents (1,4-dioxane; methanol; 10 mM sodium phosphate buffer (PB) pH 7; or PB, pH 7 containing 0.5% DMSO, v/v). Absorption spectra of these samples were taken at room temperature (RT) using a Hewlett-Packard 8452A diode array spectrophotometer equipped with the Olis Globalworks software. Emission and excitation spectra were recorded at RT using a Jobin Yvon Horiba FluoroMax-3 spectrofluorometer. Fluorescence spectra were collected using a 2 × 10 mm fluorescence cuvette, positioned for the light to pass through the shorter path. Emission spectra (405 nm excitation) were corrected by subtracting the solvent’s fluorescence followed by normalization to account for any differential absorption of TFCH and TFCZ at the excitation wavelength.

### Cell culture

A549 cells were grown in F12K media (ATCC); MDCK cells were grown in DMEM:F-12 (ATCC); and LLC-PK1 cells were grown in DMEM with 4.5 g/L glucose, L-glutamine, and sodium pyruvate (Corning). All standard cell culture media were supplemented with 10% FBS (ThermoFisher Scientific, Sigma or Atlanta Biologicals) and 1% antibiotic-antimycotic (ThermoFisher Scientific). All serum**-**free media (SFM) were supplemented only with 1% antibiotic-antimycotic. Cells were maintained in a mammalian cell culture incubator. LLC-PK1 cells were grown for 72 h and MDCK cells were grown overnight on collagen**-**coated coverglasses or 96-well plates prior to treatment.

### LOD calculation

Different concentrations of TFCZ in DMSO were added to A549 cell lysate. The final concentration of DMSO in all samples was 0.5% (v/v). The samples were excited at 405 nm and emission was recorded using a Synergy Mx microplate reader (BioTek) at 525 nm. The limit of detection (LOD) of TFCZ in the cell lysate was obtained by plotting the fluorescence of TFCZ as a function of concentration and calculated as described elsewhere [2].

### Influx of TFCH

A549 cells were imaged before and after < 5 min of adding TFCH to the cells. A 405 nm laser was used for excitation and a Zeiss confocal microscope was used for imaging.

### Washing out TFCH

A549 cells were incubated for 5 min in standard media containing 0.5% DMSO (v/v) with or without 20 µM TFCH. The cells were either lysed immediately or washed before lysis. The washing step involved discarding the TFCH-containing media, two 3 min incubation periods with fresh media, and one rinse with PBS (10 mM sodium phosphate buffered saline, pH 7.4). Emission spectra were obtained by exciting the lysates at 372 nm.

### Platereader assay by TFCH

A549 cells were grown in standard media or SFM for ~24 h. Media were then discarded and the respective media containing 1, 2, 5, or 10 µM TFCH in 0.5% DMSO (v/v) were added and incubated for 1.5 h. The cells were rinsed once with Dulbecco’s phosphate buffered saline (DPBS) and the fluorescence of the samples in DPBS were read in a Synergy Mx microplate reader (BioTek). The samples were excited at 405 nm and the emission was measured at 525 nm.

The cells were then subjected to a sulforhodamine B (SRB) assay, as described before [1]. Optical density (OD) was recorded at 570 nm (OD_570_), which was used to represent cell density. The fluorescence intensity recorded at 525 nm (F_525_), representing TFCH-associated carbonyls, was then normalized to account for variable cell densities by taking the ratio of F_525_ and OD_570_. The background signal generated by the samples treated with vehicle (0.5% DMSO, v/v) was subtracted from the signal of those treated with TFCH. Two independent experiments, each with six intra-experimental replicates, were performed. The average of the control samples treated with 1 µM TFCH was set to 100 in each independent experiment. The graph shows the mean ± SEM of the fluorescence of the sample (mean of 12 wells) relative to the average of the control treated with 1 µM TFCH.

The fold increase in fluorescence was calculated as the ratio of signal generated in the serum starved samples and the unstarved control samples. The data in the corresponding table are reported as mean ± SEM of experimental replicates.

### TFCH-mediated detection and visualization of cellular carbonyls generated by different models of oxidative stress

#### SFM-induced

A549 cells were grown in standard media or SFM for 24 h followed by incubation with 20 µM TFCH in 0.5% DMSO for 30 min. The cells were washed as previously described and imaged live at room temperature within 20 min after fluorophore incubation using a Zeiss confocal microscope. Ex: 405 nm; Em: LP 420.

In another set, control (cells grown in standard media) or serum-starved (grown in SFM) A549 cells were incubated with 2 µM TFCH in 0.5% DMSO for 30 min and imaged immediately, without washing out excess fluorophore.

In a different set, the control and the serum starved cells (A549, MDCK, LLC-PK1) were allowed to react with 20 µM TFCH for 40-60 min. The cells were then rinsed with PBS and fixed with 4% paraformaldehyde in PBS (PFA solution) for 15 min. The cells were washed twice more with PBS, mounted with ProLong^TM^ Gold antifade mountant (ThermoFisher Scientific), and cured overnight. The cells were imaged using a confocal microscope.

#### Hydrogen peroxide-induced

MDCK or LLC-PK1 cells were treated with vehicle (water) or 400 µM hydrogen peroxide (H_2_O_2_) for 2 h. The media was replaced with fresh media containing 20 µM TFCH in 0.5% DMSO (v/v). After a 30 min incubation, the cells were washed, fixed, and imaged as described previously.

Menadione-induced

MDCK or LLC-PK1 cells were treated with vehicle (DMSO, 0.05% v/v) or 100 µM menadione for 1 h prior to the addition of 20 µM TFCH in 0.5% DMSO (v/v). After 30 min, the cells were washed, fixed, mounted, and imaged as described previously.

#### Gentamicin and Cisplatin-induced

LLC-PK1 and MDCK cells grown on collagen-coated coverglasses were treated with 0.15 μg/mL of cisplatin (*cis*-Diamineplatinum(II) dichloride, Sigma) or 0.58 mg/mL of gentamicin (gentamicin sulfate salt, Sigma) for 24 h. A final concentration of 20 µM TFCH in 0.5% (v/v) DMSO was added to the samples and incubated for 40-60 min before washing, fixing and imaging. Cells that did not receive any drug treatment were used as a control.

### CellROX Green-mediated detection of reactive oxygen species (ROS) in different models of oxidative stress

All treatment methods employed for preparing the samples for subsequent staining with CellROX Green (ThermoFisher Scientific) were the same as that of TFCH. The CellROX Green staining was performed according to the manufacturer’s instructions. Briefly, the cells were treated with 5 μM CellROX Green for 30 min, washed, and fixed (as described previously) before imaging.

### Image acquisition, quantification and analysis for cell-based assays using TFCH and CellROX Green:

Cells stained with TFCH were excited using the 405 nm laser and the emission was monitored using LP420 or between 420-700 nm. Cells stained with CellROX Green were excited using the 488 nm laser and emission was monitored using LP505.

All images were obtained using a Zeiss or a Leica confocal microscope. For visual clarity, the photomicrographs were processed using identical parameters for each cell line within an experimental set.

Thresholded photomicrographs were used for quantification of signal/cell (IntDen/cell) using Fiji/ImageJ (NIH). Data are reported as mean ± SEM of signal per cell relative to the average of the control. The average of the control in each experimental set was set to 100. Data from a minimum of 4 experimental sets, with a minimum of 30 cells per sample per set, are reported in the graph. Where indicated, a two-tailed unpaired t-test with Welch’s correction or a one-way ANOVA with Dunnett’s multiple comparisons test was performed using GraphPad to compare each treatment with the control. The percent difference in mean and standard error was also calculated using GraphPad. **p*<0.05, ***p*<0.01, ****p*<0.001, *****p*<0.0001, and not significant (ns) was *p*>0.05.

### Resazurin-based assay

Cisplatin or gentamicin drug stocks were prepared fresh in appropriate cell culture media prior to each experiment. LLC-PK1 or MDCK cells grown on collagen-coated 96-well plates (100 μL media/well) were treated with only media (control), cisplatin or gentamicin by adding an additional 100 μL of fresh media supplemented with or without drugs. In parallel, a separate set of wells that did not contain cells were similarly treated for background subtraction. After 20-22 h, the cell viability assay was initiated using the *In vitro* toxicology assay kit, Resazurin based (Sigma-Aldrich) following the manufacturer’s instruction. In brief, 22 μL of the reagent was added to each well and incubated for 4 h. The samples were excited at 560 nm and emission was recorded at 590 nm using a Cytation 5 imaging reader (BioTek). Emission of the treated wells without cells was subtracted from the emission of those containing cells. Each experimental set consisted of control cells (no drug treatment), and treatment with cisplatin or gentamicin. Each graph represents drug-induced decrease in cell viability relative to the control (without drug treatment). The average of the control in each experimental set was set to 100%. Error bars show SEM. Where indicated, an unpaired t-test with Welch’s correction was performed and the percent difference in mean and standard error was calculated using GraphPad

### SRB-based assay

The drug treatment was performed as described in the resazurin-based assay section. After 24 h incubation with different drug concentrations, a SRB-based assay was performed as described before with minor modifications [1]. Experimental layout and data presentation are the same as that of the resazurin-based assay. Where indicated, an unpaired t-test with Welch’s correction was performed and the percent difference in mean and standard error was calculated using GraphPad.

### Actin and ZO-1

LLC-PK1 and MDCK cells grown on collagen-coated coverglasses were treated with different concentrations of drugs. After 24 h, the samples were processed for immunocytochemistry to assess the status of ZO-1 and actin. A polyclonal ZO-1 antibody (ThermoFisher Scientific), followed by AlexaFlour 488 goat anti-rabbit IgG (H+L), was used to stain ZO-1. Subsequently, the cells were stained with Rhodamine Phalloidin and Hoechst 33342 to probe for F-actin and the nucleus, respectively. Cells were imaged using a Zeiss confocal microscope. The intensity of actin-stress fibers within each cell (IntDen/cell) was quantified by Fiji/ImageJ (NIH) in thresholded images. The intensity (IntDen) of cell membrane-associated ZO-1 was quantified by selecting a fixed region of interest drawn with the “rectangle” tool. Emission channels representing actin and ZO-1 staining are presented in the figure. For visual clarity, the photomicrographs were processed using identical parameters for each cell line within an experimental set. Graphs represent the intensity of actin stress fiber per cell or ZO-1 intensity, both relative to the control. The average of the control was set to 100. An unpaired t-test with Welch’s correction was performed and the percent difference in mean and standard error was calculated using GraphPad.

### Detection of carbonyls in rat tissue slices

Male Spraque-Dawley rats (250 g) purchased from Charles River Laboratories were utilized. Rat kidney slices were prepared and maintained as described elsewhere with minor modifications [3]. After perfusion using PBS, the kidneys were sliced, washed, and equilibrated in HBSS pH 7.4 with 5% CO2/95% O2 at 37 ᵒC. Subsequently, the kidney tissue slices were treated with different concentrations of cisplatin (37.5, 75, 150 μg/ mL) or gentamicin (1.2, 2.3, 4.6 mg/mL) in HBSS buffer. After 1 h, TFCH (final concentration: 2 μM) was added to each sample and incubated for 30 min at 37 °C. Multiple control samples were processed in parallel. To assess the basal level of carbonyls in the kidney slices, untreated (no drug) samples incubated with TFCH (final concentration: 2 μM) were used. To ensure that the fluorescence detected was not contributed by residual drug itself, tissue samples were exposed to the highest concentrations of cisplatin or gentamicin used and treated with DMSO (vehicle of TFCH, 0.5%, v/v) for 30 min. At the end of the fluorophore or vehicle treatment, all samples were washed with PBS and incubated with a periodate-lysine paraformaldehyde (PLP) fixative for 20 min. The samples were washed 4-5 times with PBS before mounting with ProLong^TM^ Gold antifade mountant (ThermoFisher Scientific) and cured overnight. The samples were imaged using a Zeiss confocal microscope. Representative photomicrographs were assembled (stitched) from multiple sections of the renal cortex. For visual clarity, the photomicrographs presented in a set were processed using identical parameters unless otherwise mentioned.

Thresholded photomicrographs of untreated (control), cisplatin (150 μg/ mL), or gentamicin (4.6 mg/mL) treated kidney slices were used for quantification of renal tubule-associated fluorescence. The fluorescence intensity (IntDen) of the renal tubule was quantified by selecting a fixed region of interest drawn with the “rectangle” tool in Fiji/ImageJ software. Data are reported as mean ± SD of signal relative to the average of the control. The average of the control was set to 100. A two-tailed unpaired t-test with Welch’s correction was performed using GraphPad for statistical analysis. Based on this analysis, *****p*<0.0001.

Animal experiments were approved by the institutional committee on Research Animal Care, in accordance with the National Institutes of Health guide for the care and use of the laboratory animals.
